# SaccuFlow – A High-throughput Analysis Platform to Investigate Bacterial Cell Wall Interactions

**DOI:** 10.1101/2021.04.14.439837

**Authors:** Alexis J. Apostolos, Noel J. Ferraro, Brianna E. Dalesandro, Marcos M. Pires

## Abstract

Bacterial cell walls are formidable barriers that protect bacterial cells against external insults and oppose internal turgor pressure. While cell wall composition is variable across species, peptidoglycan is the principal component of all cell walls. Peptidoglycan is a mesh-like scaffold composed of crosslinked strands that can be heavily decorated with anchored proteins. The biosynthesis and remodeling of peptidoglycan must be tightly regulated by cells because disruption to this biomacromolecule is lethal. This essentiality is exploited by the human innate immune system in resisting colonization and by a number of clinically relevant antibiotics that target peptidoglycan biosynthesis. Evaluation of molecules or proteins that interact with peptidoglycan can be a complicated and, typically, qualitative effort. We have developed a novel assay platform (SaccuFlow) that preserves the native structure of bacterial peptidoglycan and is compatible with high-throughput flow cytometry analysis. We show that the assay is facile and versatile as demonstrated by its compatibility with sacculi from Gram-positive bacteria, Gram-negative bacteria, and mycobacteria. Finally, we highlight the utility of this assay to assess the activity of sortase A from *Staphylococcus aureus* against potential anti-virulence agents.

## Introduction

Bacterial cell walls have many functions but none more important than acting as a protective barrier from potentially harsh external elements and attacks from host defense mechanisms.^1, 2^ Bacteria can be categorized based on the type of cell wall they possess and there are distinctive features of each that can pose a challenge, or opportunity, for the host the immune system or antibiotics.^3, 4^ In the case of Gram-positive bacteria, their cell wall is composed of a thick peptidoglycan (PG) sacculus layer followed by a cytoplasmic membrane. PG is a mesh-like polymer made up of repeating disaccharide units of *N*-acetylglucosamine (GlcNAc) and *N*-acetylmuramic acid (MurNAc). Attached to each MurNAc unit is a short peptide, referred to as the stem peptide, that ranges from 3 to 5 amino acids in length. The stem pentapeptide sequence is typically L-Ala-*iso*-D-Glu-L-Lys (or *meso*-diaminopimelic acid [*m*-DAP])-D-Ala-D-Ala, although there are considerable variations in the primary sequence.^5^

Crosslinking can occur between neighboring stem peptides and the degree to which the PG is crosslinked can influence the cell wall rigidity and integrity.**Error! Reference source not found**. The thick PG layer in Gram-positive bacteria is generally believed to be a permeability barrier to molecules in the extracellular space.^6–9^ While most small molecules can potentially sieve through the PG, there is evidence that the PG is a formidable barrier for the permeation of larger molecules.^10–12^ This feature has significant implications for molecules with intracellular and cell membrane targets; one example being the membrane attack complex (MAC), a product of the complement system that targets bacterial cell membranes. The MAC is prevented from reaching the cytoplasmic membrane of Gram-positive bacteria due in part to its inability to penetrate through the PG layer.^13, 14^ Gram-positive pathogens can also alter the composition of their PG to become resistant to specific classes of antibiotics. For example, both *Staphylococcus aureus* (*S. aureus*) and *Enterococcus faecium* can alter the PG structure to resist the antibacterial actions of vancomycin.^15–17^ Due to its critical role in bacterial cell survival, PG is often the target of antibiotics and components of the human immune system.^4, 18, 19^ Clinically and industrially important antibacterial agents (e.g., β-lactams, teixobactin, vancomycin, bacitracin, and moenomycin) target components of the PG biosynthesis as integral parts of their mechanism of actions. Similarly, components of the human immune system (e.g., lysozyme, LysM-displaying proteins, and PG recognition proteins [PGRPs]) must reach the PG to impart their response onto the invading bacterial pathogen.

Despite the importance of characterizing interactions with the PG scaffold of bacteria, there are limited methods to investigate the specific binding of molecules to PG. More specifically, there is a clear need for a method to determine PG binding interactions that (1) retains the natural composition of native PG, (2) is applicable to all types of bacteria, (3) preserves the polymeric nature of native PG, (4) is readily attainable in high yields, and (5) is compatible with quantitative, high-throughput analysis platforms. There are several challenges with obtaining PG samples for analysis. The use of synthetic PG mimics provides some advantages such as sample homogeneity and the ability to install specific changes to the PG primary sequence. In fact, a number of important contributions to the field have been made using synthetic PG mimics.^20–24^ Our own group has extensively used synthetic PG mimics in studying cell wall biosynthesis and remodeling. ^25, 26^ However, there are some severe drawbacks to using synthetic analogs, including the lack of commercially available building blocks for the disaccharide units and *m*-DAP. In addition, there is potential for the loss of polyvalent interactions, which are prevalent in PG-binding molecules, using monomeric PG analogs. Alternatively, PG fragments can be obtained by digesting isolated native PG (sacculi) with a hydrolase enzyme and performing chromatography.^27, 28^ This last-resort method is fraught with challenges due to the difficulty in the separation of fragments, convoluted characterization of fragments (e.g., site of amidation), and low yields. Finally, the use of intact cells to study PG interactions has advantages in terms of retaining the polymeric nature and preservation of the native composition (even if variable across the PG scaffold). However, isolation of the effect of PG interaction alone in the background of surface bound proteins and other potential biomacromolecules is a major hurdle. Herein, we report a new method (SaccuFlow) that uses sacculi isolated directly from Gram-positive, Gram-negative, or mycobacterial organisms in combination with flow cytometry to assay PG binding interactions. To the best of our knowledge, flow cytometry has not yet been systematically used to evaluate isolated bacterial PG and/or its interactions with binding partners.

## Results and Discussion

We reasoned that murein sacculus from bacteria, which is a single macromolecular scaffold made of PG, would provide a superior platform to analyze PG interactions due to its ease of isolation and proper mimicry of the PG composition and structure. While sacculi analysis has been implemented for decades, these studies have focused primarily on mass spectrometry analysis of digested low molecular weight PG fragments^29–31^ and low-throughput qualitative analyses techniques (e.g., confocal microscopy^32^ and pull-down assays^33^). We envisioned a that a higher-throughput and quantitative sacculus analysis platform could be achieved by performing the analysis using flow cytometry. Sacculi should be readily detected *via* flow cytometry because its size resembles that of the bacterial cell.^34^ Significantly, sacculi preparations are routinely performed with relative ease from almost any type of bacteria with these isolation steps.

Initially, we set out to benchmark the detection of bacterial sacculi from a Gram-positive organism using a standard flow cytometer. We chose to site selectively tag the sacculi with a fluorescent handle by metabolic labeling of the PG scaffold. Our research group^35–39^, and others^26, 35, 37, 38, 40–47^, have previously demonstrated that co-incubation of unnatural D-amino acids, including those modified with fluorophores, metabolically label PG of live bacteria cells. More specifically, fluorescein conjugated D-Lys (**D-LysFl**) from the culture media is expected to replace the 5^th^ position D-alanine of the PG stem peptide of *S. aureus* during the biosynthesis and remodeling of new PG (**Figure 1A**). Generally, 1-2 % of the stem peptides in the PG scaffold of *S. aureus* is labeled when cells are treated overnight with the D-amino acid label under the assay conditions. Bacterial sacculi were isolated using standard procedures from whole cells, which were expected to yield fluorescently labeled sacculi (**Figure 1B**). Our results revealed that labeled sacculi formed a tight population of events that could readily be distinguished from background debris (**Figure 1C**). Most significantly, fluorescence levels of sacculi from cells treated with **D-LysFl** were 17-fold higher than cells treated with the control amino acid **L-LysFl**. Unlike **D-LysFl**, its enantiomer does not become incorporated as part of the bacterial PG scaffold. The same sacculi was imaged using confocal microscopy, thus confirming that the sample analyzed was primarily fluorescent sacculi (**Figure S1**).

**Figure 1.**
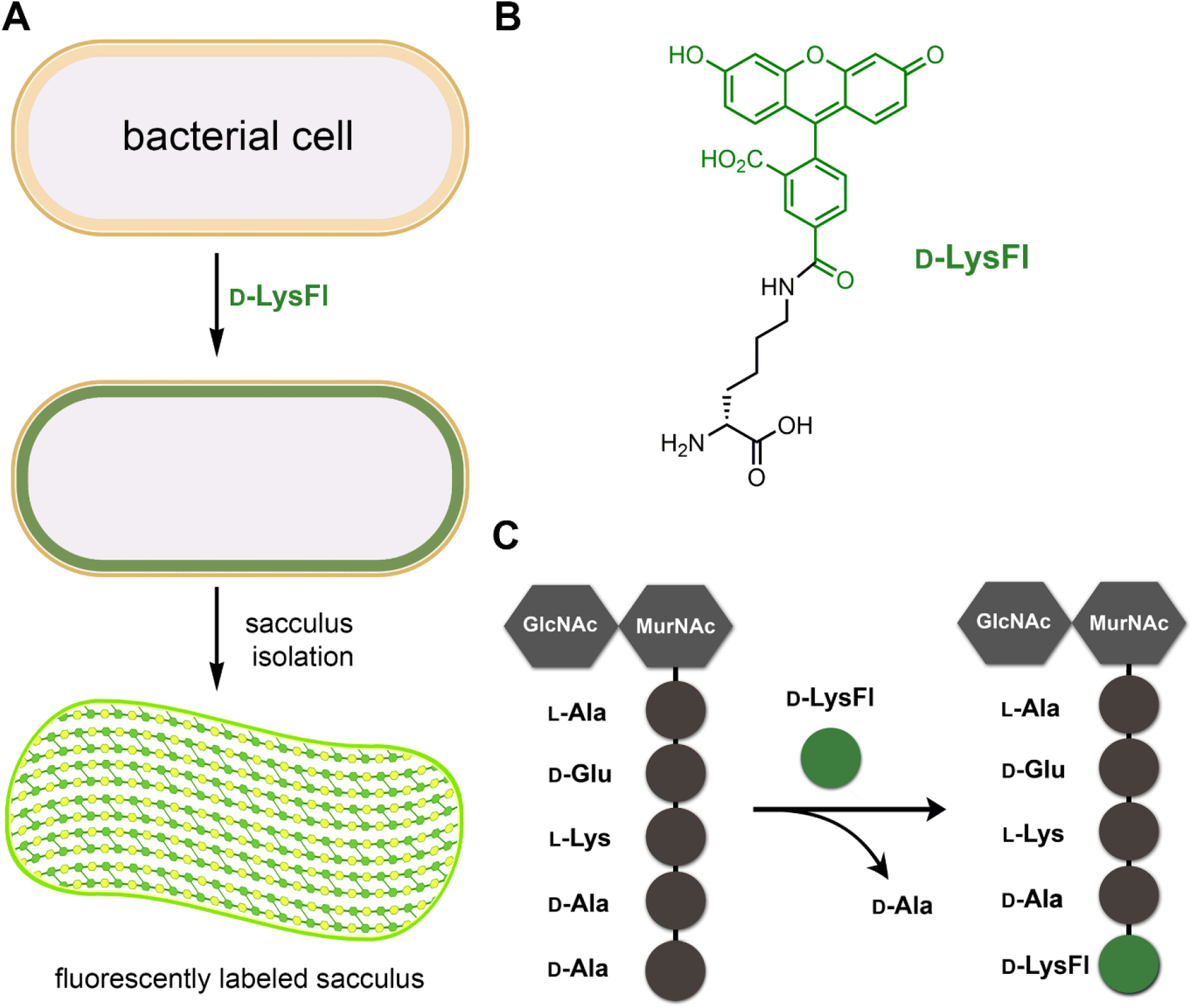
(**A**) Assay workflow of SaccuFlow for the labeling of sacculi. (**B**) Chemical structure of **D-LysFl**. (**C**) Cartoon representation of the metabolic swapping of the terminal D-ala from the bacterial PG with the exogenously supplied **D-LysFl**.

**Figure 2.**
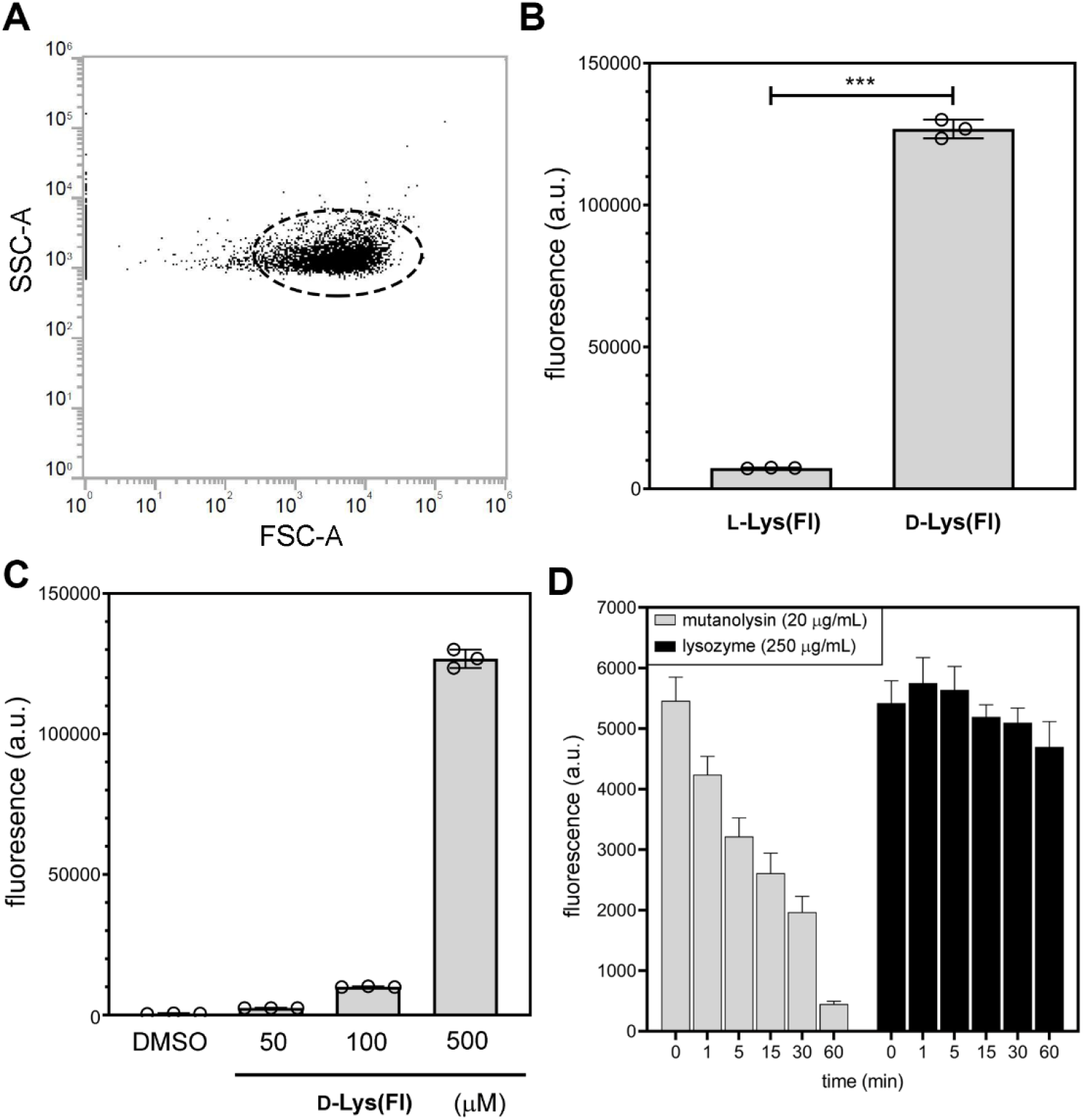
(**A**) Side and forward scatter plot of *S. aureus* sacculi. (**B**) Flow cytometry analysis of sacculi isolated from *S. aureus* (ATCC 25923) treated overnight with 500 μM of **D-LysFl** or **L-LysFl**. (**C**) Flow cytometry analysis of sacculi isolated from *S. aureus* (ATCC 25923) treated overnight with varying concentrations of **D-LysFl**. (**D**) Flow cytometry analysis of sacculi isolated from *S. aureus* (ATCC 25923) treated overnight with 500 μM of **D-LysFl** and incubated with either mutanolysin or lysozyme. Data are represented as mean +/− SD (n = 3). *P*-values were determined by a two-tailed *t*-test (* denotes a *p*-value < 0.05, ** < 0.01, ***<0.001, ns = not significant).

Next, a series of additional experiments were performed to confirm that the events being detected on the flow cytometer were, indeed, *S. aureus* sacculi. Sacculi labeled with **D-LysFl** were subjected to treatment with two PG hydrolases: lysozyme and mutanolysin (**Figure 1D**). As expected, treatment with mutanolysin resulted in a time-dependent reduction in fluorescence levels due to the release of PG fragments. *O*-acetylation of *S. aureus* PG renders it insensitive to lysozyme digestion and, likewise, lysozyme treatment.^48^ Digestion by mutanolysin was also found to be concentration dependent (**Figure S2**). To show the versatility of the tagged sacculi in a flow cytometry platform, we prepared sacculi from organisms that had been co-incubated with either **D-** or **L-LysAz** (**Figure S3**,**4**). We anticipated that **D-LysAz** treated cells would provide an orthogonal click handle on the PG scaffold that could be covalently linked to a variety of compounds containing the corresponding reactive moieties. Subsequent treatment with cyclooctynes should result in strain-promoted alkyne-azide cycloaddition (SPAAC) ligation with azide groups on the sacculi. ^49, 50^ For this assay, azide-tagged sacculi isolated from *S. aureus* were incubated with dibenzocyclooctyne (DBCO) linked to fluorescein.

Three different linker lengths composed of polyethylene glycol spacers were tested to investigate how size of the molecule can impact permeation across the sacculi (**Figure 3A,B**). Linkage of **Dfl0** monitored by fluorescence was ~8 fold over the DMSO vehicle. Additionally, corresponding with what we have previously seen on whole *S. aureus* cells, we observed a decrease in fluorescence as the length of the spacer increased with **Dfl3** and **Dfl9** (**Figure 3C**). Strikingly, after chemical removal of the wall teichoic acids (WTA), an increase in fluorescence was observed with the shortest linker length. WTA is an anionic polymer that is covalently anchored onto PG of some Gram-positive organisms.^51–54^

**Figure 3.**
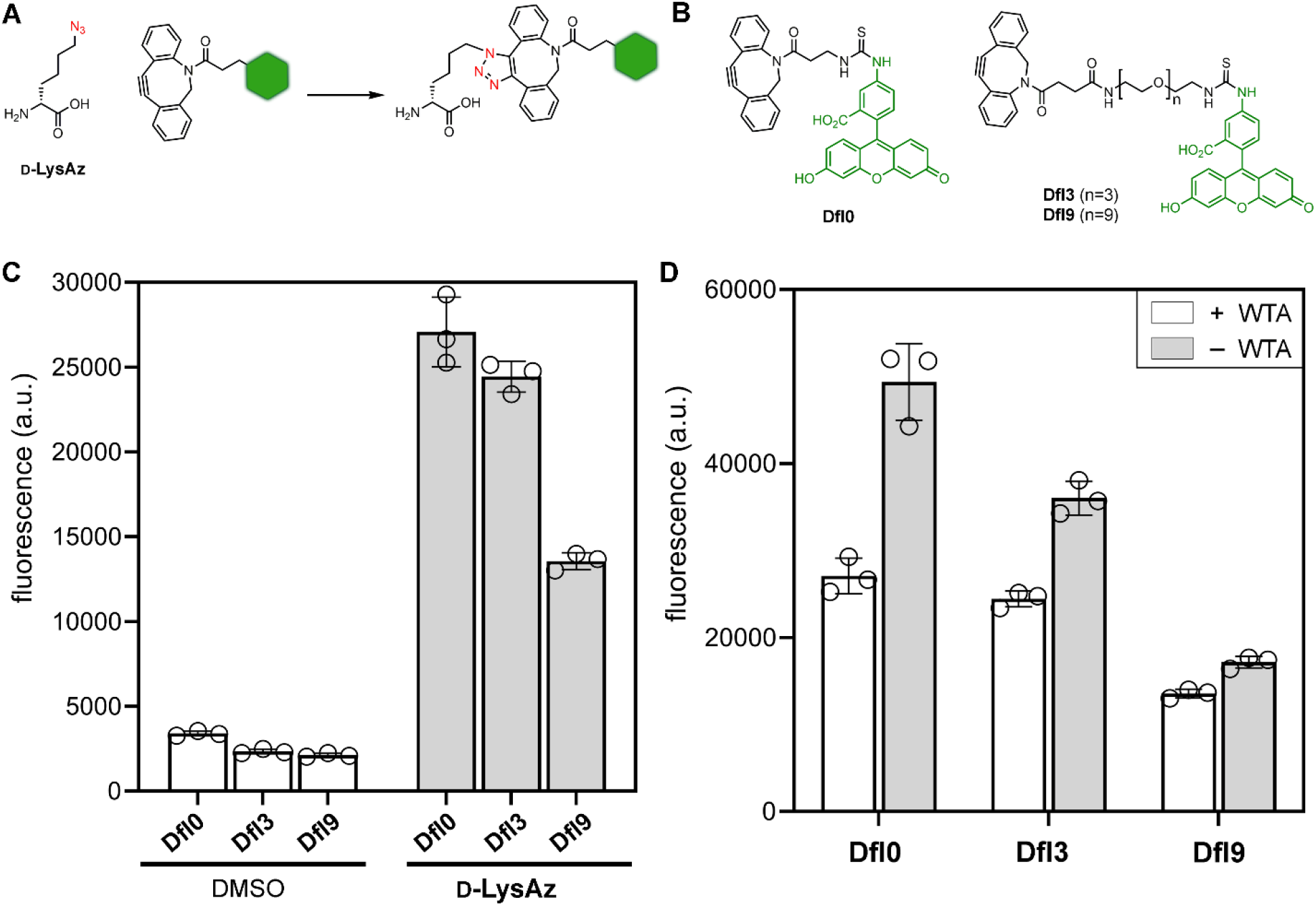
(**A**) Chemical structures representing the SPAAC reaction between azide and a strained alkyne in DBCO. (**B**) Chemical structures of **Dfl0, Dfl3**, and **Dfl9**. (**C**) Flow cytometry analysis of sacculi isolated from *S. aureus* (ATCC 25923) treated overnight with 1 mM of **D-LysAz** or DMSO. Subsequently, sacculi were incubated with 25 μM of **Dfl0, Dfl3**, or **Dfl9**. (**D**) Flow cytometry analysis of sacculi isolated from *S. aureus* (ATCC 25923) treated overnight with 1 mM of **D-LysAz**. Where noted, sacculi were chemically treated to remove WTA. Subsequently, sacculi were incubated with 25 μM of **Dfl0, Dfl3**, or **Dfl9**. Data are represented as mean +/− SD (n = 3). *P*-values were determined by a two-tailed *t*-test (* denotes a *p*-value < 0.05, ** < 0.01, ***<0.001, ns = not significant).

It is well established that WTA can block or impede interaction of extracellular molecules with the PG scaffold. These results highlight the role that surface biopolymers play in accessibility to the PG scaffold and further demonstrates the ability to use flow cytometry to systematically assess this important feature related to bacterial cell wall biology. Similarly, to test the applicability of this method to mycobacterial and Gram-negative organisms, *Mycobacterium smegmatis* (*M. smegmatis*) and *Escherichia coli* (*E. coli*) were grown in media supplemented with either **D-** or **L-LysAz** and their sacculi were harvested following the standard procedures for each classification. The isolated PG was then treated with **Dfl0** and analyzed by flow cytometry, showing that only sacculi from cells that had been treated with **D-LysAz** labeled extensively with **Dfl0** (**Figure S4**). This highlights the adaptability of SaccuFlow to bacterial species of varied classifications.

Having shown that SaccuFlow was operational and facile, we then set out to demonstrate that it could report on features related to PG-binding antibiotics. Vancomycin, a glycopeptide antibiotic that specifically hydrogen-binds the D-Ala-D-Ala fragment of the stem peptide, conjugated to BODIPY (**VBD**)^55^ was analyzed for binding using SaccuFlow. As anticipated, *S. aureus* sacculi demonstrated ~8-fold BODIPY fluorescence over background, suggesting that vancomycin bound to isolated PG was detectable by flow cytometry (**Figure 4A**). To show that fluorescence levels were reflective of specific binding interactions, bacterial sacculi was treated with a synthetic analog of the PG, L-Lys-D-Ala-D-Ala. As expected, increasing concentrations of L-Lys-D-Ala-D-Ala led to decreasing fluorescence levels associated with the decrease in binding events of **VBD** to sacculi (**Figure 4B**). To show the versatility of this assay amongst other strains, we isolated *Bacillus subtilis* (*B. subtilis*) sacculi from a wildtype (WT) strain and from a strain lacking *dacA* (*dacA*Δ), the gene responsible for the D-alanyl-D-alanine carboxypeptidase.^56^ **VBD** would be expected to bind *dacA*Δ *B. subtilis* sacculi to a higher level in comparison to WT, as lack of the carboxypeptidase would present additional D-Ala-D-Ala binding points for **VBD**. Our results demonstrate a ~**3**-fold increase in binding of **VBD** to *dacA*Δ sacculi as compared to WT, as monitored by SaccuFlow (**Figure 4C**). Next, sacculi samples were isolated from an *Enterococci faecium* (*E. faecium*) strain that has a vancomycin inducible resistant phenotype based on activation of the membrane receptor VanS.^57, 58^ The outcome of this activation is the intracellular removal of the terminal D-alanine in the PG precursor, which should reduce the bindings available to **VBD**. Consistent with this phenotypic switch, analysis of VRE sacculi showed that there was a significant decrease in fluorescence associated with VRE pre-treated with vancomycin (**Figure 4D**). Together, these results confirm that SaccuFlow can be readily adopted to monitor and quantify interactions of PG-binding molecules, which can have applicability in studying drug resistance or establish therapeutic interventions.

**Figure 4.**
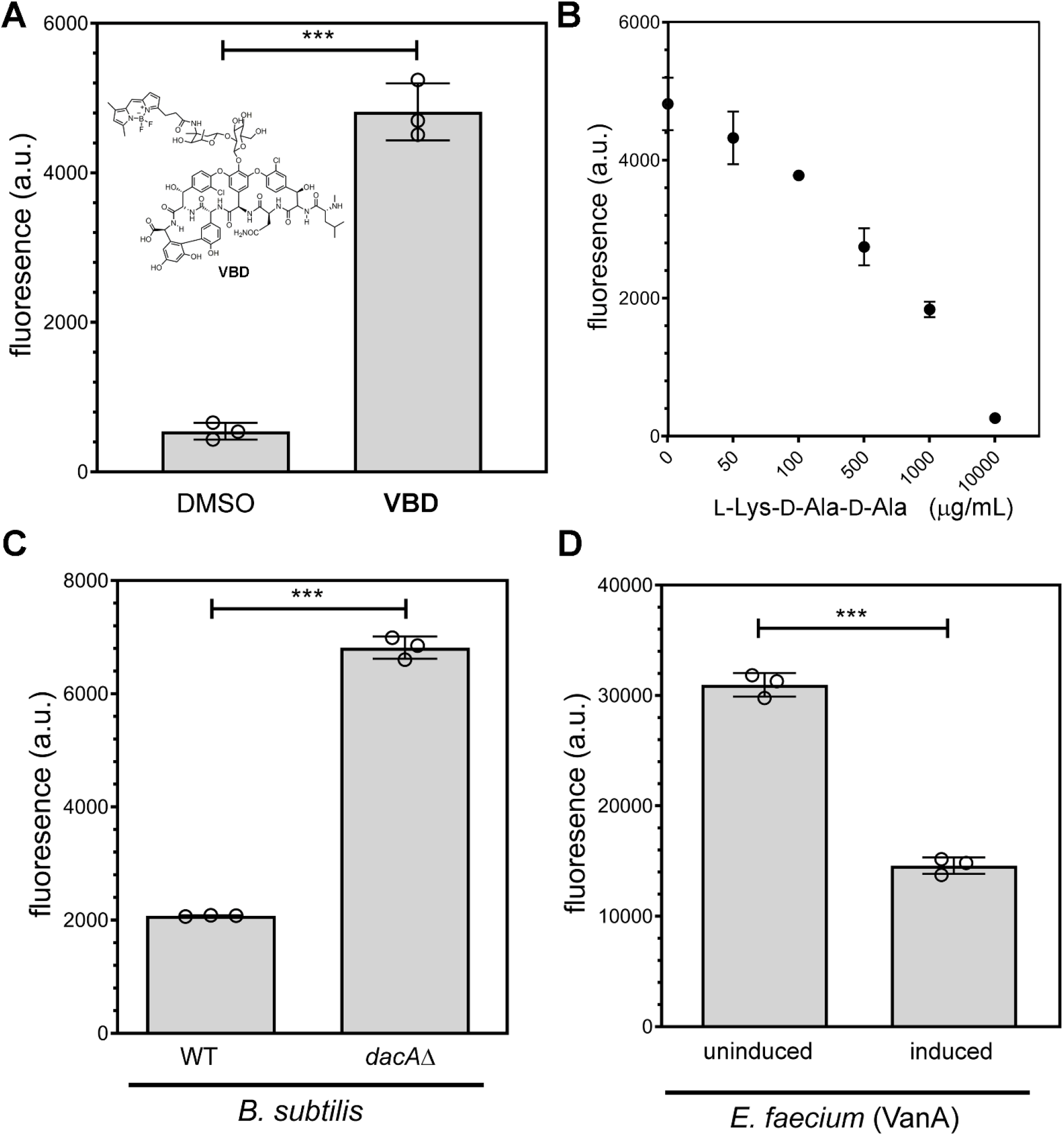
(**A**) Flow cytometry analysis of sacculi isolated from *S. aureus* (ATCC 25923). Subsequently, sacculi were treated with **VBD** (4 μg/mL) or DMSO. (**B**) Flow cytometry analysis of sacculi isolated from *S. aureus* (ATCC 25923). Subsequently, sacculi were treated with **VBD** (2 μg/mL) and increasing concentrations of L-Lys-D-Ala-D-Ala. (**C**) Flow cytometry analysis of sacculi isolated from *B. subtilis* (WT and *dacA⊿*). Subsequently, sacculi were treated with **VBD** (2 μg/mL). (**D**) Flow cytometry analysis of sacculi isolated from *E. faecium* vanA (with and without pre-incubation of vancomycin). Subsequently, sacculi were treated with **VBD** (4 μg/mL). Data are represented as mean +/− SD (n = 3). *P*-values were determined by a two-tailed *t*-test (* denotes a *p*-value < 0.05, ** < 0.01, ***<0.001, ns = not significant).

We tested the ability of the SaccuFlow assay to evaluate the processing and remodeling of PG. More specifically, we evaluated the possibility of tracking the activity of the enzyme sortase A from *S. aureus*. Sortase A is a transpeptidase that recognizes the sorting sequence LPXTG (where X is any amino acid) and thereby anchors proteins that contain this sequence onto the bacterial PG.^59, 60^ *S. aureus* utilizes sortase A to heavily decorate the surface of the PG with proteins capable of improving host colonization and interfering with human immune response. Therefore, sortase A is considered an attractive drug target and small molecules that inhibit sortase A could prove to be promising anti-virulence agents. Prior efforts to investigate sortase A activity have generally used minimal peptide mimics of PG (e.g., penta-glycine), which may not be entirely representative of the features of the PG (or its lipid anchored precursor). For our assay, sacculi isolated from *S. aureus* were incubated with purified sortase A originating from *S. aureus* and a sorting signal substrate modified with a fluorophore. We monitored the fluorescence of the sacculi over time *via* flow cytometry and, as expected, observed an increase in fluorescence as sortase A linked the fluorescent sorting signal substrate to the isolated PG (**Figure 5A**). Further, sacculi treated with methanethiosulfonate (MTSET), a covalent inhibitor of sortase^61^, or no sortase at all, demonstrated fluorescence that was at background levels (**Figure 5A**). These results provide clear demonstration that SaccuFlow can be utilized to investigate processing of PG in a quantitative manner.

**Figure 5.**
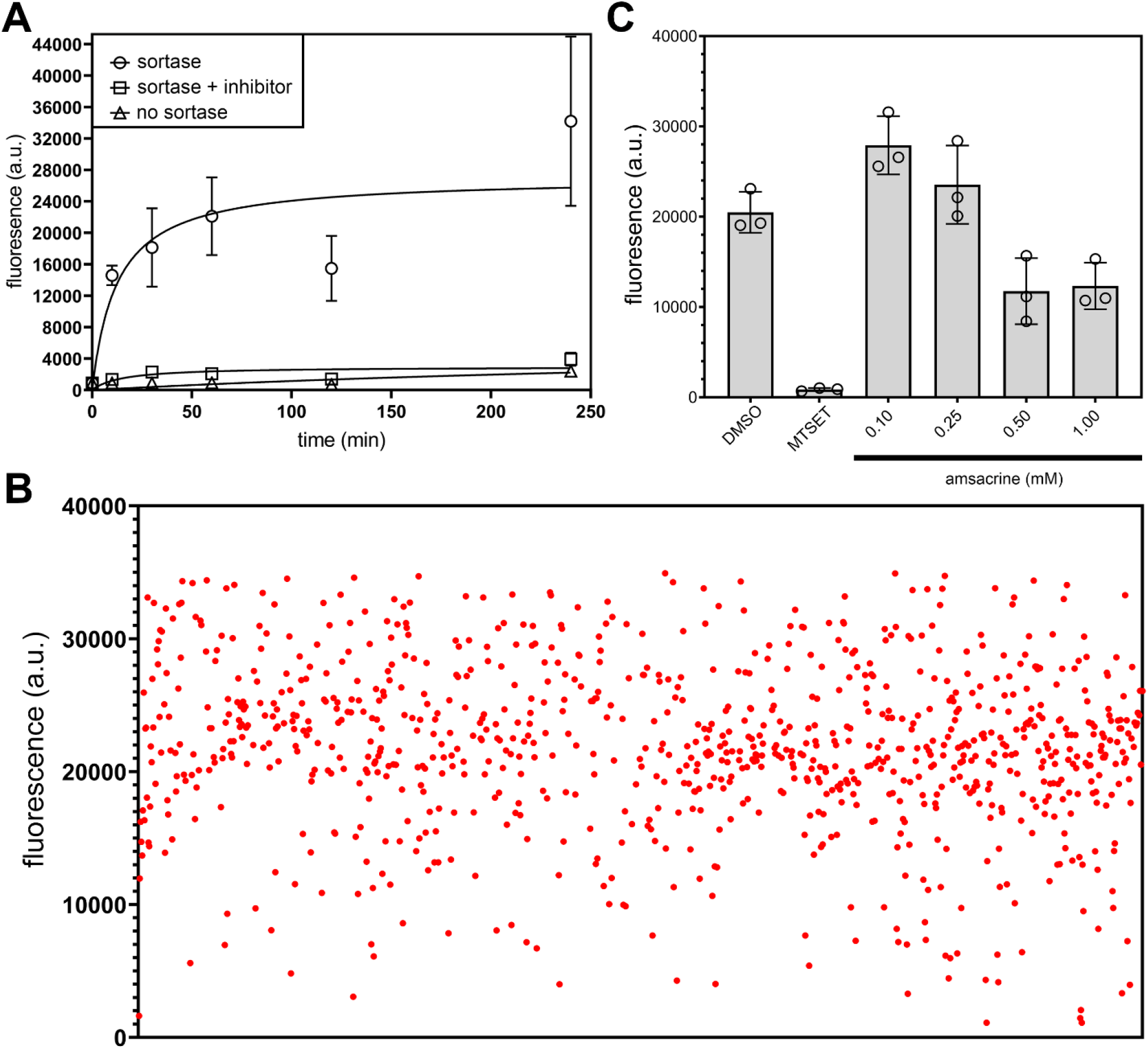
(**A**) Time course analysis of sortase activity using isolated sacculi from *S. aureus* (ATCC 25923) in the presence of the fluorescent sorting signal substrate (100 μM). Flow cytometry analysis of sacculi treated with sortase, the covalent inhibitor MTSET (1 mM), and in the absence of sortase. (**B**) Screening of LOPAC 1280 library using 384-well format. Individual dots represent members of the library in the presence of isolated sacculi and sorting signal substrate (100 μM). (**C**) Flow cytometry analysis of sortase activity in the presence of designated conditions. Data are represented as mean +/− SD (n = 3).

Finally, to demonstrate the high-throughput capability of SaccuFlow, we used the established parameters of the sortase A assay to determine the ability of a library of 1,280 pharmacologically active compounds to inhibit sortase A activity (LOPAC 1280 library).

Using isolated *S. aureus* sacculi, sortase A, and the f sorting signal substrate, we monitored the fluorescence readout of the sacculi against individual members of the compound library. We predicted that a reduction in fluorescence would correspond to a reduction of sortase A activity. Our screening results revealed a total of 18 compounds as potential inhibitors of sortase (**Figure 5B**). Of interest, amsacrine, which was previously reported to inhibit Mycobacterial topoisomerases^62^ as well as D-alanylation of teichoic acids in *S. aureus*^63^, was identified as a potential inhibitor. We further tested the ability of amsacrine to inhibit sortase A in a titration assay, which showed a reduction in fluorescence levels in a concentration-dependent manner (**Figure 5C**). This pilot screen demonstrated the feasibility of SaccuFlow to be miniaturized and used in a high-throughput screen. Moreover, these results highlight the potential of SaccuFlow to assess essential enzyme operations within the PG scaffold and identify substrates or inhibitors of those enzymes of interest, paving the way to the discovery of potential therapeutics against these validated drug targets.

## Conclusion

In conclusion, we have described the implementation of a new flow cytometry assay (SaccuFlow) that makes use of the mechanical strength and native composition of bacterial sacculi. By adopting the assay to flow cytometry, we showed that it is possible to gain quantitative information on interactions of molecules with PG. Tagging of bacterial sacculi with orthogonal epitopes led to the demonstration of the sieving process through the PG scaffold based on molecular size. Moreover, versatility of the assay platform was also demonstrated by analyzing sacculi from three different species of Gram-positive bacteria, one Gram-negative bacterium, and a mycobacterium. Binding studies with fluorescently labeled vancomycin showed that SaccuFlow can reveal PG interaction dynamics, including phenotypic changes due to drug resistance. Finally, enzymatic processing of PG by sortase A was performed to highlight the versatility of this assay in studying PG biology. Given the important nature of PG binding molecules, including several clinically relevant antibiotics and components of the innate immune system, we expect that this assay platform will find wide usage across microbiology studies.

