## Supporting Information for "SaccuFlow – A High-throughput Analysis Platform to Investigate Bacterial Cell Wall Interactions"

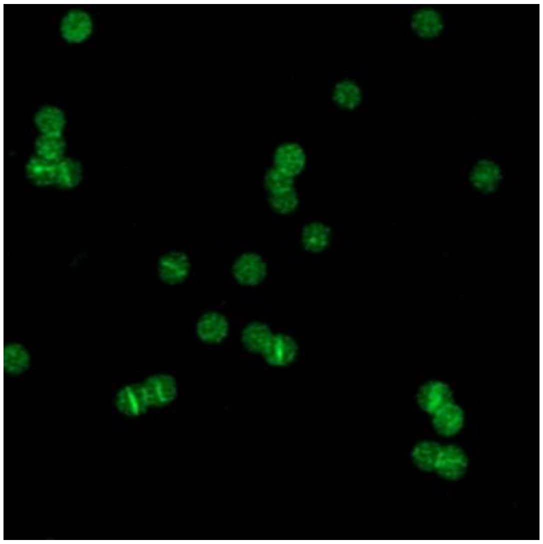

**Supplemental Figure 1.** Confocal microscopy of sacculi isolated from *S. aureus* (ATCC 25923) that was treated with 100 μM D-**LysFl** overnight.

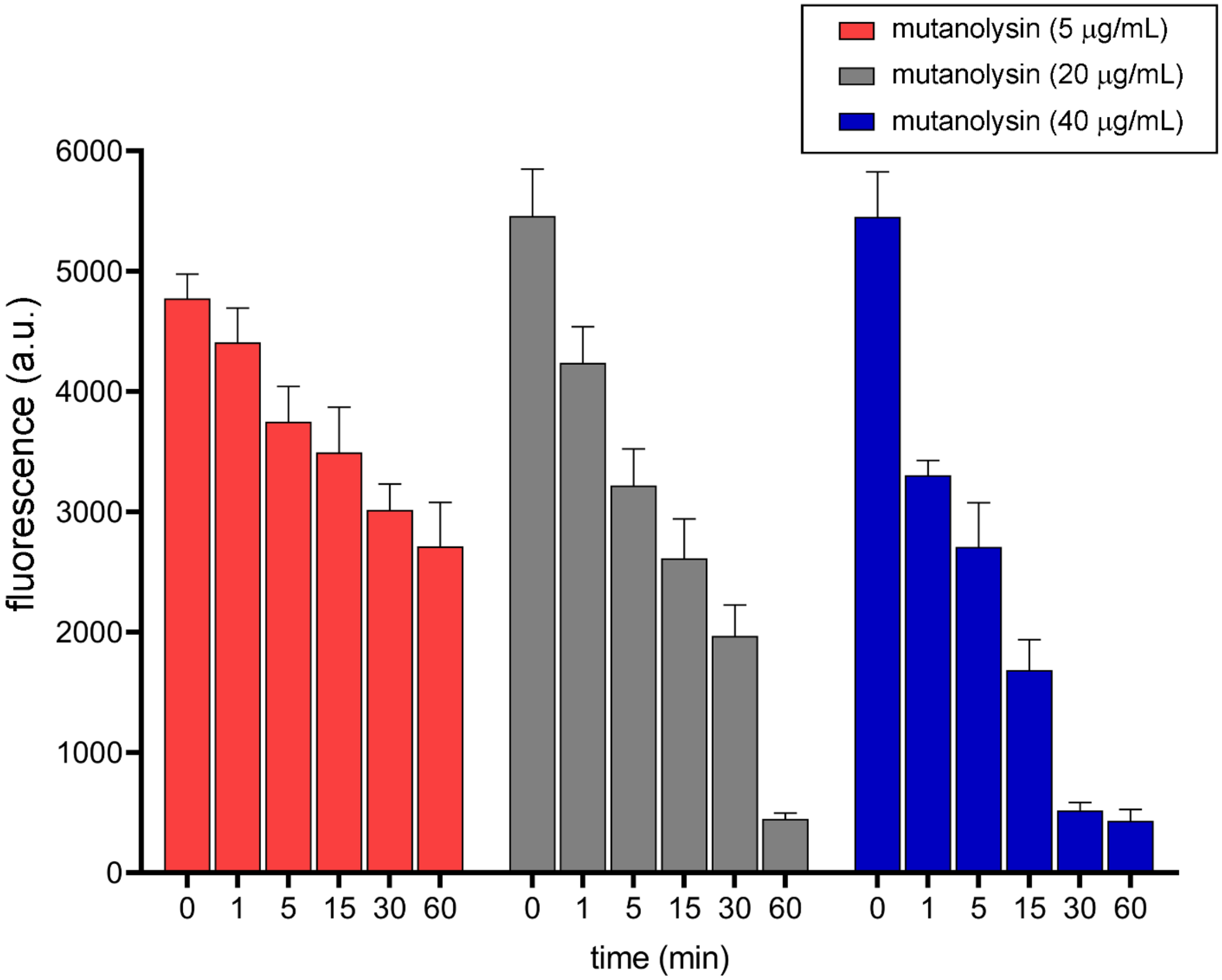

**Supplemental Figure 2.** Fluorescence read out of sacculi isolated from *S. aureus* (ATCC 25923) that was treated with 100 µM D-**LysFl** overnight and subjected to degradation by increasing concentrations of mutanolysin overtime.

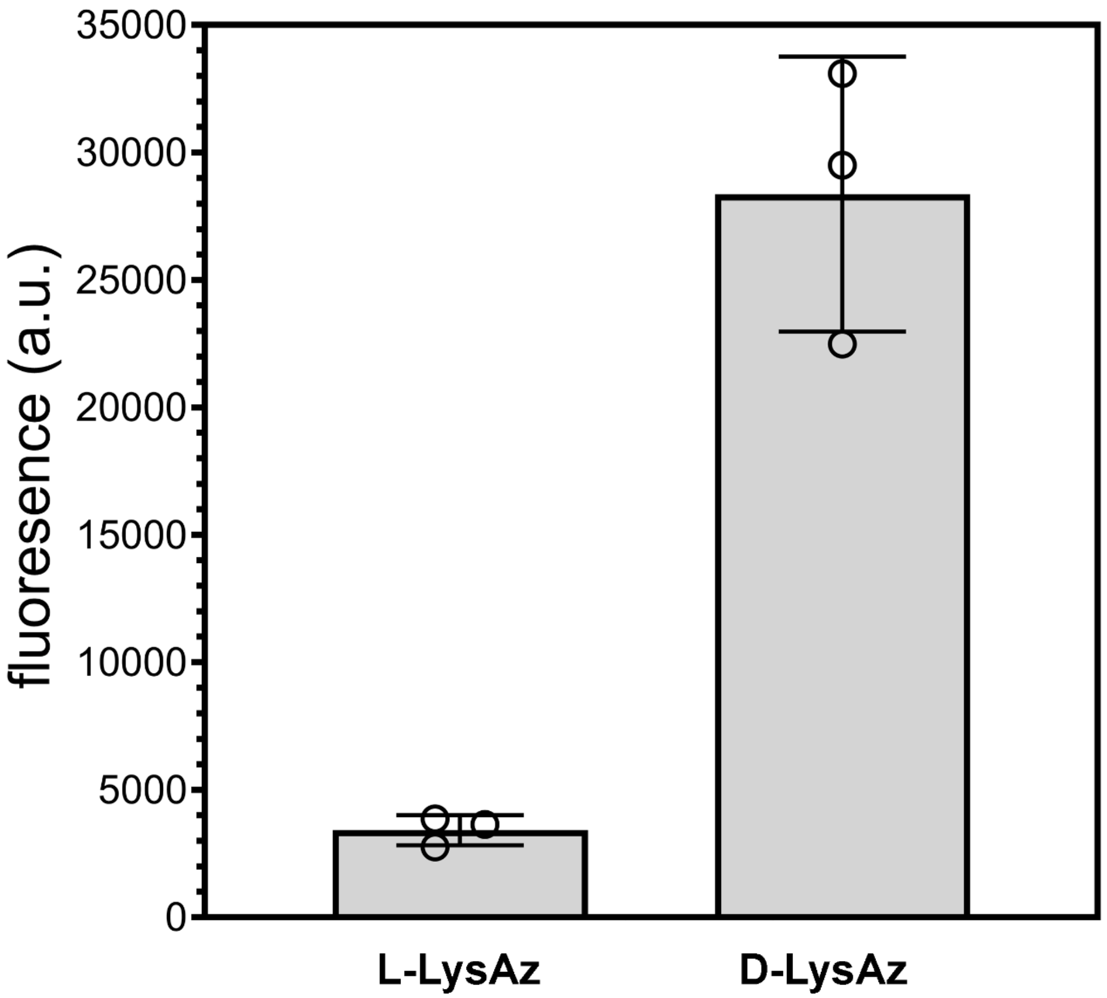


**Supplemental Figure 3.** Fluorescence read-out of sacculi isolated from *S. aureus* (ATCC 25923) that was incubated with 1 mM **L-** or D-**LysAz** overnight, and subsequently treated with 25 µM DBCO-FITC.

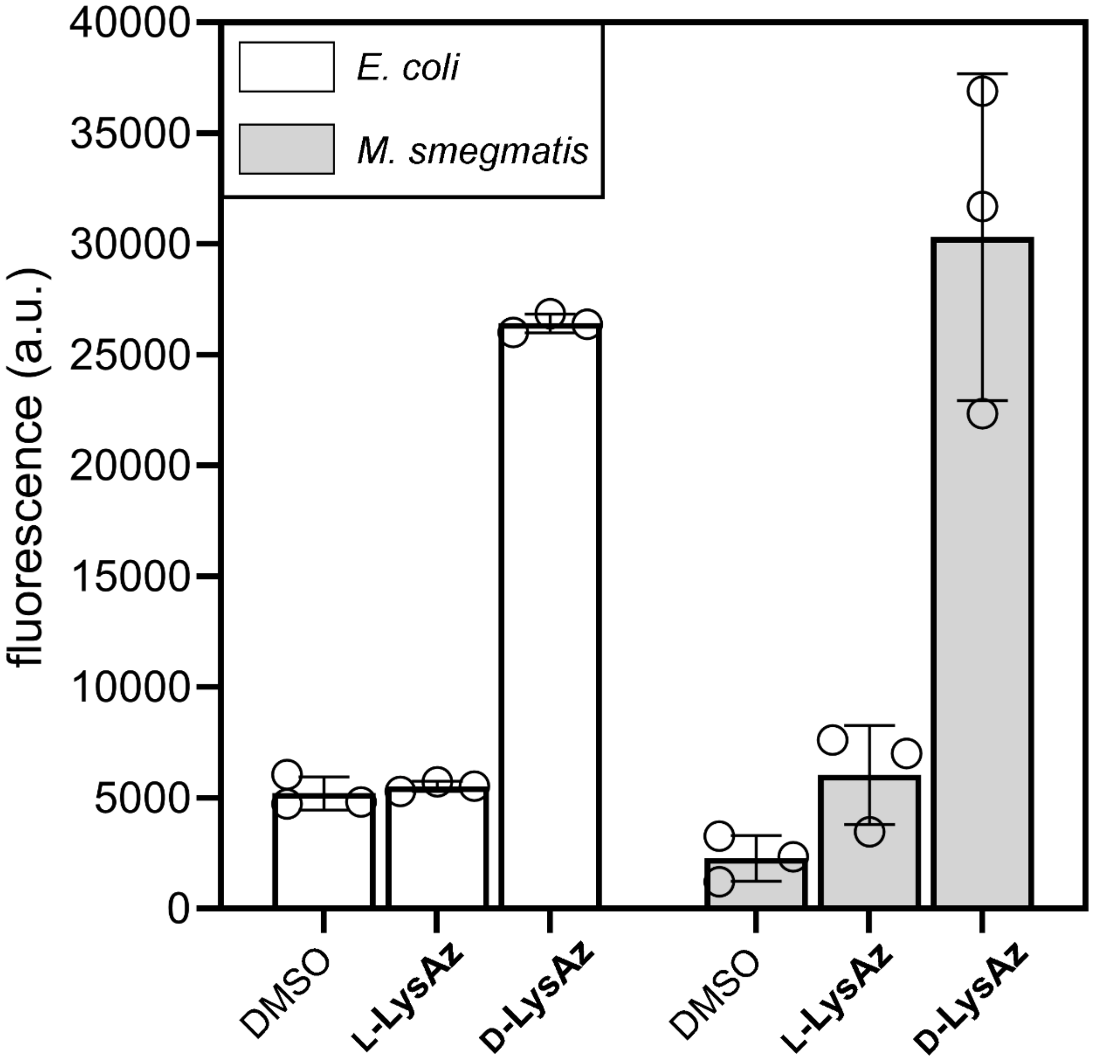


**Supplemental Figure 4.** Fluorescence read-out of sacculi isolated from *E. coli* BW25113 or *M. smegatis* WT that was incubated with 1 mM **L-** or D-**LysAz** overnight, and subsequently treated with 25 µM DBCO-FITC.

**Materials.** All peptide related reagents (resin, coupling reagent, deprotection reagent, amino acids, and cleavage reagents) were purchased from ChemImpex or BroadPharm. Bacterial strains *Staphylococcus aureus* (*S. aureus*), *Bacillus subtilis* (*B. subtilis*), *B. subtilis dacA*Δ, and *Escherichia coli* (*E. coli*) were grown in lysogeny broth, *Enterococcus faecium* (*E. faecium*) was grown in tryptic soy broth, *Mycobacterium smegmatis* (*M. smegmatis*) were grown in lysogeny broth supplemented with 0.05% Tween 80 for all experiments.

**Peptidoglycan Isolation of *S. aureus*.** Lysogeny broth (LB, 25 mL) containing 1 mM D- or L-Lys-azido, 50, 100, or 500 µM D-Lys(FITC), 500 µM L-Lys(FITC), or no supplemented probe was prepared. *S. aureus* bacteria were added to the LB medium (1:100) and allowed to grow overnight at 37°C with shaking at 250 rpm. The cells were harvested and then resuspended in 1X PBS, boiled for 25 min, and then centrifuged at 14,000g for 15 min at 4°C. Cells were then placed in 15 mL of 5% (w/v) sodium dodecyl sulfate (SDS) and boiled for 25 min followed by centrifugation at 14,000g for 15 min at 4°C. Following centrifugation, cells were boiled again in 25 mL of 4% (w/v) SDS for 15 min followed by centrifugation using same parameters as before. Cells were then washed 6 times with 60 °C DI water to remove all SDS. After washing, cells were resuspended in 6 mL of 20 mM Tris buffer (pH 8.0). Pellets were treated with 800 µg DNase for 24 h followed by trypsin (800 µg) for another 24 h (37°C shaking at 115 rpm). Pellets were boiled for 25 min followed by centrifugation at 14,000g for 15 min at 4°C, resuspended in 2 mL of 1X PBS, and further diluted for analysis by flow cytometry. In samples where wall teichoic acid removal was desired the pellets after the last centrifugation step were resuspended 1 M HCl and incubated for 4 h at 37°C with shaking. The pellet was harvested by centrifugation for 10 min at 4,000 rpm and washed with dd$H_2O$ until the pH of the supernatant reached 6-7. The final pellet was resuspended in 1X PBS, and further diluted for analysis by flow cytometry with an Attune NxT flow cytometer, equipped with a 488 nm laser and 525/40 nm bandpass filter. The data were analyzed using the Attune NXT Software.

**Peptidoglycan Isolation of *B. subtilis* and *B. subtilis dac*Δ.** LB was supplemented with 1X kanamycin during the growth of the *B. subtilis dacA*Δ*.* The rest of the isolation protocol follows that of the *S. aureus* isolation protocol detailed above.

**Peptidoglycan Isolation of *E. faecium.*** Tryptic soy broth (TSB) was supplemented with 16 µg/mL of vancomycin in order to induce the resistance phenotype during the growth of the *E. faecium.* The rest of the isolation protocol follows that of the *S. aureus* isolation protocol detailed above.

**Peptidoglycan Isolation of *E. coli.*** LB (100 mL) containing 1 mM D- or L-Lys-azido, or no supplemented probe was prepared. *E. coli* BW25113 bacteria were added to the LB medium (1:100) and allowed to grow overnight at 37°C with shaking at 250 rpm. The cells were chilled on ice, then harvested at 8,000g for 10 min at 4°C. The resulting pellet was resuspended in 3 mL of 1% (w/v) NaCl, then added slowly to 6 mL of 8% SDS that was in a boiling water bath with a stir bar. Samples were boiled for 5 h, followed by incubation at room temperature with continued stirring overnight. The following morning, samples were boiled for 2 h and then pelleted at 100,000g for 10 min at 25°C. Resulting pellets were suspended in 3 mL of 4% SDS, boiled for 2 h with stirring. Samples were collected using the same parameters and washed 5 times with sterile DI $H_2O$ to remove all SDS. Pellets were resuspended in 1 mL of 1X PBS and 200 µg/mL of trypsin was added followed by incubation at 37°C with shaking at 115 rpm for 2 h. A second dose of trypsin was added, and samples were incubated overnight. The following morning, one-tenth volume of 10% SDS was added and samples were placed in an oil bath at 100°C for 2 h. Sacculi were collected as before, and washed 3X with sterile DI $H_2O$ to remove residual SDS. The final pellet was lyophilized and resuspended in 400 µL of $H_2O$ for further analysis by flow cytometry.

**Peptidoglycan Isolation of *M. smegmatis.*** LB (100 mL) with 0.05% Tween 80 containing 1 mM D- or L-Lys-azido, or no supplemented probe was prepared. *M. smegmatis* bacteria were added to the LB medium (1:100) and allowed to grow overnight at 37°C with shaking at 250 rpm. Cells were then harvested at 14,000g for 15 min at 4°C and washed 1X in a minimal volume of 1X PBS. Pellets were resuspended in 10 mM $NH_4HCO_3$ with a protease inhibitor cocktail (Sigma SRE005-1BO). This suspension was lysed using a tip sonicator (Fisher Scientific) at 60% amplitude for 60 seconds, with at least 60 seconds on ice in between. This was repeated for 5 cycles. The sonicate was digested with 10 µg/mL DNAse and RNase for 1 h at 4°C, then harvested at 27,000g for 30 min at 4°C. The pellet was resuspended in PBS with 2% SDS and incubated for 1 h at 50°C with constant stirring. This was collected using the same centrifugal parameters and the SDS process was repeated 2X. The resulting pellet was resuspended in PBS with 1% SDS and 0.1 mg/mL Proteinase K at 45°C for 1 h with stirring. The sample was then heated to 90°C for 1 h and collected as above. This step at 90°C was repeated 2X. The sample was washed 2X with PBS and 4X with dH2O. This yielded mycolic-arabinogalactan-peptidoglycan (mAGP). The final mAGP sample was lyophilized and resuspended in $H_2O$ for further SPAAC chemistry and analysis by flow cytometry.

**Enzymatic Degradation.** Isolated 100 µM D-Lys(FITC) labeled *S. aureus* sacculi samples were pelleted in a 96-well plate at 4,000 rpm for 4 min and then resuspended in a 1X PBS solution containing either 5, 20, or 40 µg/mL mutanolysin or 250 µg/mL lysozyme and allowed to incubate at 37°C, however a zero time point was taken before any samples were subjected to enzyme treatment. A portion of the cells were then taken at 1, 5, 15, 30, and 60 minutes. At each time point, the collected bacteria were resuspended solution of 1X PBS containing 4% formaldehyde to quench the enzymatic activity. Samples were analyzed by flow cytometry as described above.

**SPAAC Reaction with Isolated Sacculi.** Isolated sacculi samples (50 µL) were pelleted in a 96-well plate at 4,000 rpm for 4 min and then resuspended in a 1X PBS solution containing 25 µM DBCO-FITC. Plates were incubated at 37°C for 30 min and washed 3X with 1X PBS by centrifugation as before to remove excess FITC. Samples were then diluted 10-fold for analysis by flow cytometry as described above.

**Vancomycin-BODIPY (VBD) Binding Assays.** Isolated *S. aureus* sacculi samples were pelleted in a 96-well plate at 4,000 rpm for 4 min and then resuspended in a 1X PBS solution containing either 2 µg/mL **VBD** alone or 2 µg/mL **VBD** in conjunction with either 50, 100, 500, 1000, or 10000 µg/mLof the L-Lys-D-Ala-D-Ala peptide. The plates were incubated for 30 min at 37°C and after incubation the samples were immediately subjected to analysis by flow cytometry as described above. Isolated *B. subtilis* and *B. subtilis dacA*Δ sacculi samples were pelleted in a 96-well plate at 4,000 rpm for 4 min and then resuspended in a 1X PBS solution containing 4 µg/mL **VBD** and incubated for 30 min at 37°C. After incubation samples were washed with1X PBS and subjected to analysis by flow cytometry as described above. *E. faecium* (noninduced and induced) sacculi samples were pelleted in a 96-well plate at 4,000 rpm for 4 min and then resuspended in a 1X PBS solution containing 4 µg/mL **VBD** and incubated for 30 min at 37°C. After incubation samples were immediately subjected to analysis by flow cytometry as described above.

**Sortase A Expression and Purification**. The plasmid for sortase A from *S. aureus* was obtained from Addgene: pET28a-SrtAdelta59. Competent BL21 (DE3) *E. coli* were transformed with the plasmid. *E. coli* cells containing the plasmid from an overnight culture were diluted 1:100 and grown at 37°C until the $OD_{600}$ was 0.4-0.6. At this time, Isopropyl-β-D-thiogalactopyranoside (IPTG) was added to a concentration of 0.5 mM and the cultures were shook at 25°C for 16 h. Cells were collected at 3,000g for 30 min and resuspended in 50 mM Tris-HCl, pH 7.5, and 150 mM NaCl. Pellets were then recollected and lysed in cold lysis buffer (50 mM Tris-HCl, pH 7.5, 150 mM NaCl, 5 mM $MgCl_2$, 10 mM imidazole, 10% vol/vol glycerol, 1 mg $ml^{-1}$ DNase and 1 mg $ml^{-1}$ lysozyme). Cells were lysed using an ultrasonic cell sonicator and the lysate was separated from cellular debris by centrifugation at 20,000g for 30 min at 4°C. The supernatant was loaded onto a Ni-NTA agarose resin and eluted with 500 mM imidazole in buffer (50 mM Tris-HCl pH 7.5, 150 mM NaCl, and 10% vol/vol glycerol). The eluted protein was dialyzed against buffer without imidazole, aliquoted (protein concentration determined by Bradford assay), and stored at –20°C until use.

**Sortase A Enzymatic Assays**. Isolated *S. aureus* sacculi samples were incubated with 20 uM sortase A, 100 uM sorting signal substrate, and 1X sortase buffer (10X contains 500 mM Tris-HCl, pH 7.5, 1.5 M NaCl, 100 mM $CaCl_2$). Samples that contained the covalent inhibitor, MTSET, were run at a concentration of 1 mM. All were shook at room temperature for the given time. At each time point, 20 uL were taken out of the reaction, quenched with 0.1% TFA, and washed 3X with 100 mM Tris, 5 mM EDTA, pH 7 and freshly added 8 M urea. Samples were resuspended in a final volume of 120 uL 1X PBS and analyzed by flow cytometry as described above.

For the high-throughput scan using the LOPAC library of 1,280 small molecule compounds, the same conditions were used. However, sortase was incubated with 1 mM of the compound library and let incubate at room temperature for 20 minutes before addition of sacculi, Fl-LPMTG (sorting signal), and buffer. After the addition of the remaining reaction components, samples were shook at room temperature for 4 h before being quenched with 0.1% TFA, then washed in buffer with 8M urea as above. Samples were resuspended in a final volume of 100 uL 1X PBS and analyzed by flow cytometry as described above.

**Scheme S1. Synthesis of D-LysFl (shown) and L-LysFl**

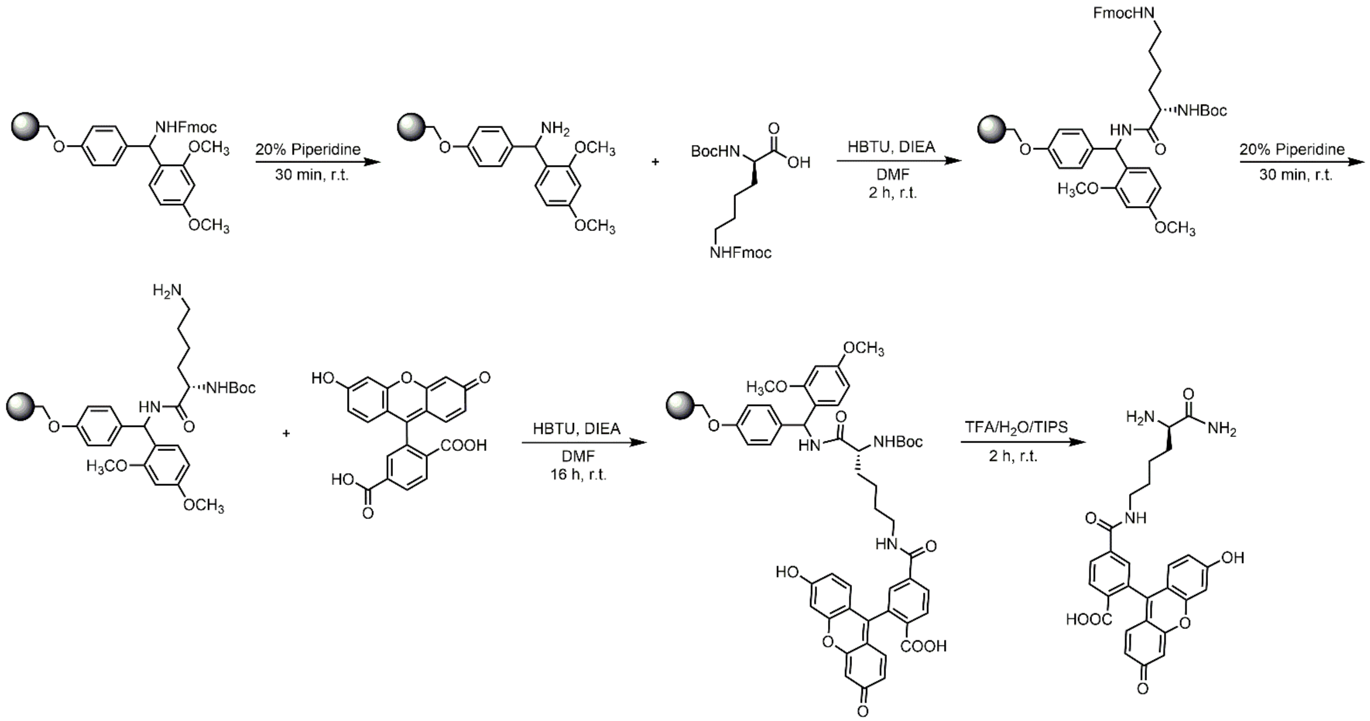


A 25 mL peptide synthesis vessel charged with rink amide resin (250 mg, 0.11 mmol) underwent the Fmoc removal procedure and was washed as described above. Boc-D-Lysine(Fmoc)-OH (5 eq, 257 mg, 0.55 mmol), HBTU (4.9 eq, 204 mg, 0.53 mmol), and DIEA (10 eq, 0.191 mL, 1.10 mmol) in DMF (15 mL) were added to the reaction vessel and agitated for 2 h at ambient temperature. After 2 h the resin was washed as previously stated and the Fmoc protecting group removal was also performed as described above followed by washing. The resin was coupled with 5,6-carboxyfluorescein (2 eq, 82 mg, 0.22 mmol), HBTU (1.9 eq, 79 mg, 0.20 mmol), and DIEA (4 eq, 0.076 mL, 0.44 mmol) in DMF (15 mL) and agitated for 16h at ambient temperature. The resin was washed as previously described and then added to a solution of TFA/$H_2O$/TIPS (95%, 2.5%, 2.5%, 20 mL) with agitation for 2 h at ambient temperature. The resin was filtered and the resulting solution was concentrated *in vacuo*. The residue was trituated with cold diethyl ether. The sample was analyzed for purity using a Waters 1525 Binary HPLC Pump using a Phenomenex Luna 5u C8(2) 100A (250 x 4.60 mm) column; gradient elution with $H_2O/CH_3CN$.

**D-LysFl**

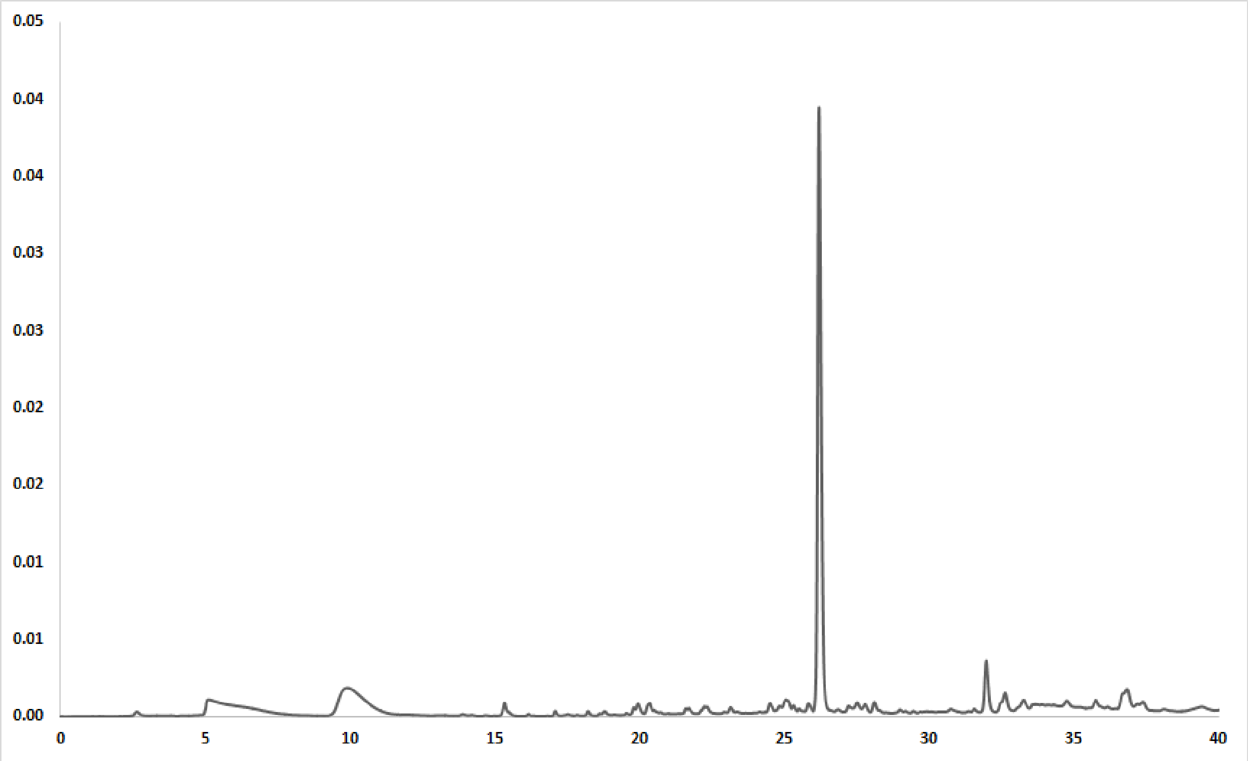


**ESI-MS calculated [M + H]$^{+}$: 504.1765, found: 504.1765**

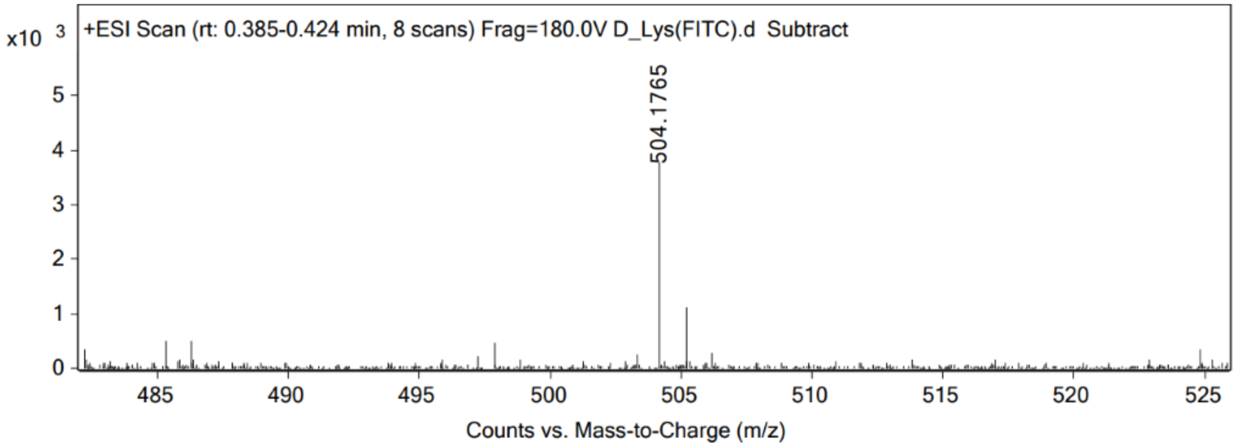


**L-LysFI**

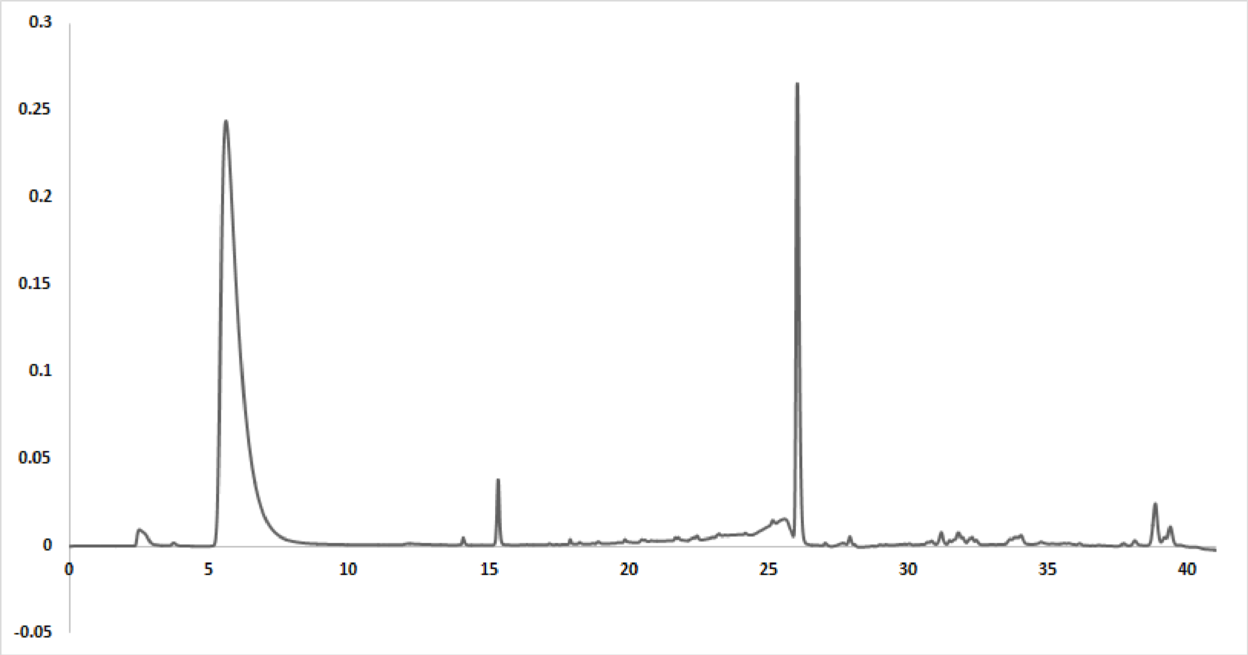


**ESI-MS calculated [M + H]$^{+}$: 504.1765, found: 504.1767**

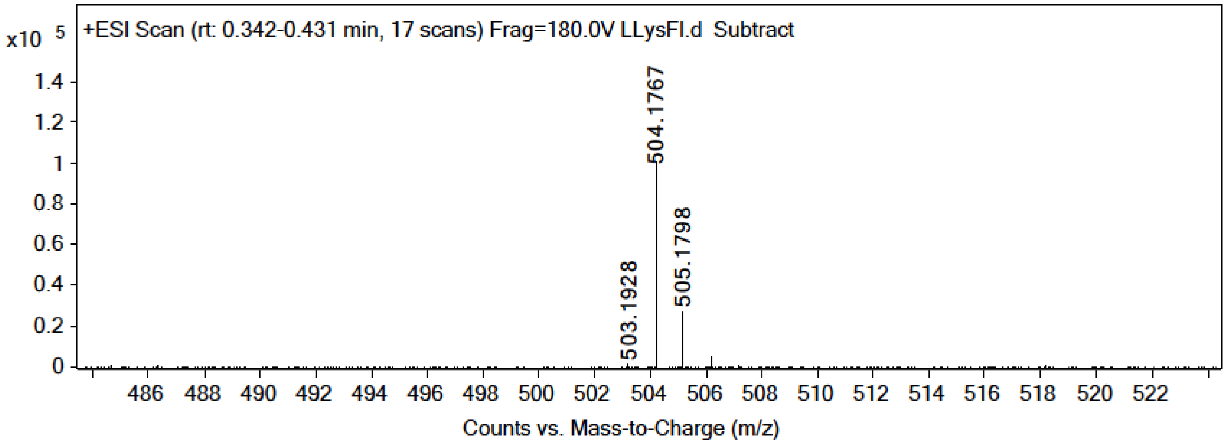


**Scheme S2. Synthesis of DBCO-peg$_9$-Fl**

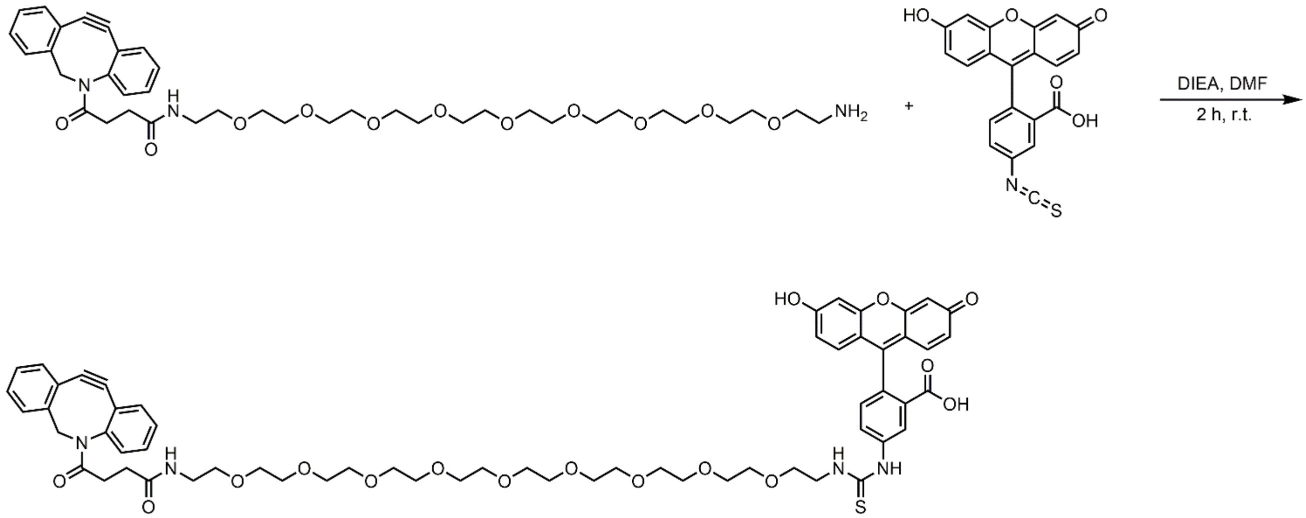


DBCO-PEG$_9$-amine was added to fluorescein isothiocyanate isomer I (2 eq) and DIEA (2 eq) in DMF and allowed to react at ambient temperature for 2 h. The reaction mix was purified using reverse phase HPLC using $H_2O$/MeOH. The sample was analyzed for purity using a Waters 1525 Binary HPLC Pump using a Phenomenex Luna 5u C8(2) 100A (250 x 4.60 mm) column; gradient elution with $H_2O$/$CH_3CN$ (DMSO signal has been subtracted).

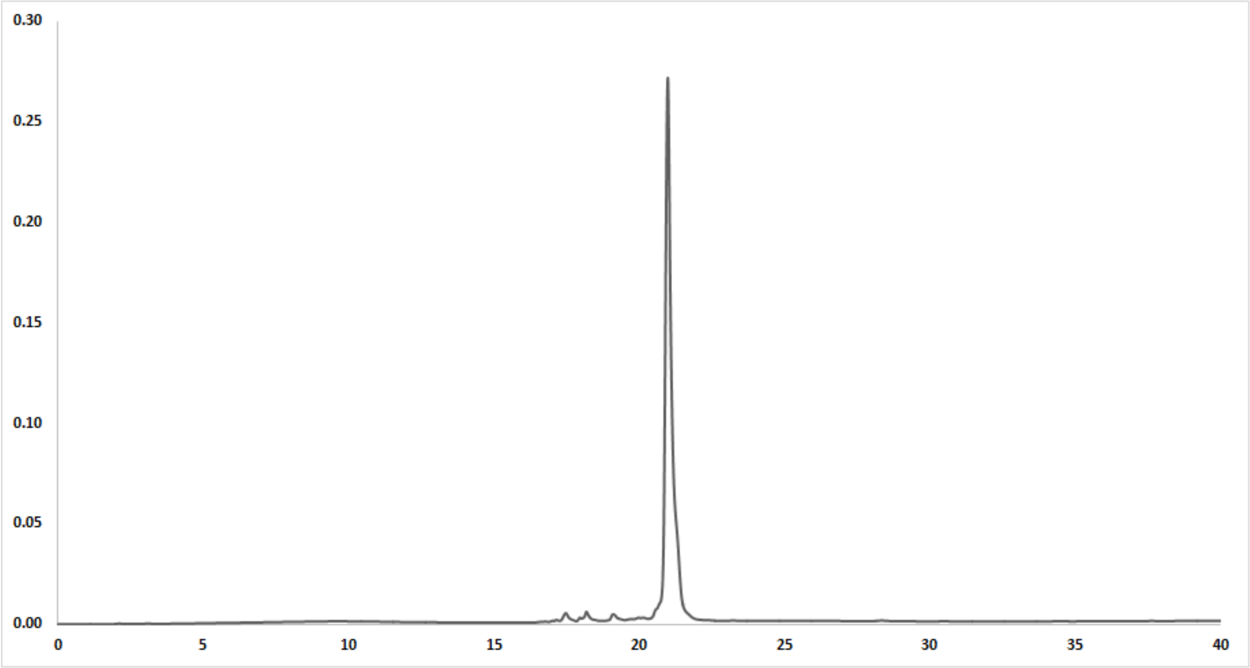

**ESI-MS calculated [M + H]$^+$: 1133.4423, found: 1133.4428**

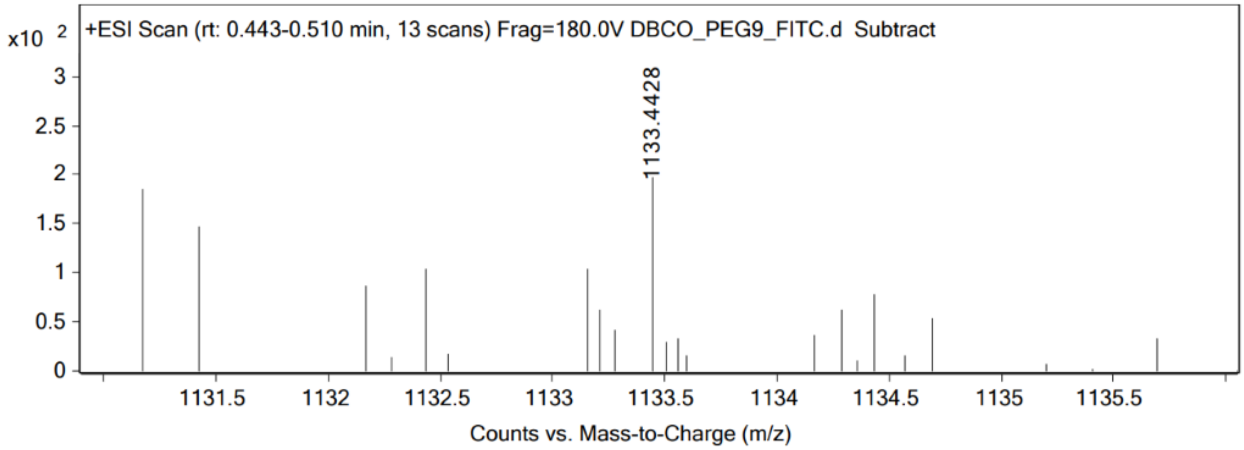


**Scheme S3. Synthesis of Fl-LPMTG**

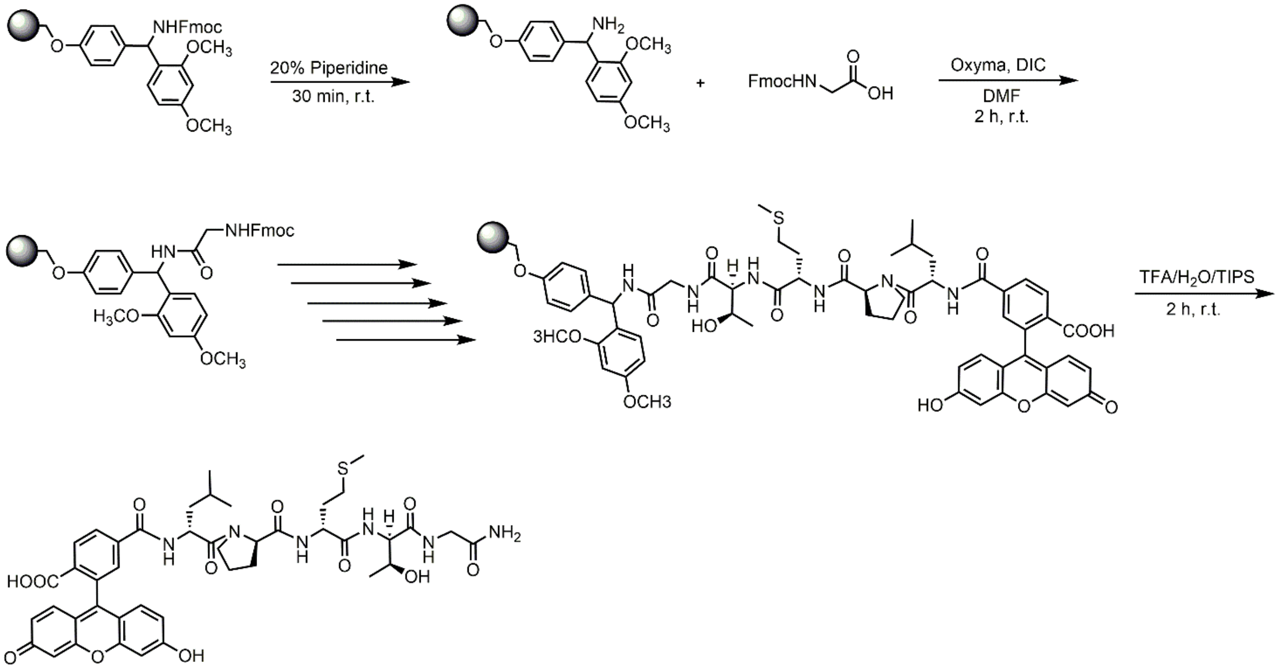


A 25 mL synthetic vessel was charged with 500 mg (0.27 mmol) of Fmoc-Rink amide resin. The Fmoc protecting group was removed with a 20% piperidine in DMF solution (15 mL) for 30 minutes at ambient temperature, then washed with MeOH and DCM (3 x 15 mL each). Fmoc-glycine-OH (3 eq, 240 mg, 0.810 mmol), Oxyma (3 eq, 115 mg, 0.810 mmol), and DIC (3 eq, 126 uL, 0.810 mmol) in DMF (15 mL) was added to the reaction vessel and agitated for 2 hours at ambient temperature and washed as previously stated. The Fmoc removal and coupling procedure was repeated as before using the same equivalencies with Fmoc-L-threonine(tBu)-OH, Fmoc-L-methionine-OH, Fmoc-L-proline-OH, and Fmoc-L-leucine-OH. The Fmoc group of L-leucine was deprotected and resin coupled with 5,6-carboxyfluorescein (2 eq, 203 mg, 0.810 mmol), HBTU (2 eq, 201 mg, 0.810 mmol), and DIEA (4 eq, 187 uL, 1.08 mmol) in DMF (15 mL) shaking over-night. The resin was washed as previously described. To remove the peptide from resin, a TFA cocktail solution (95 % TFA, 2.5% TIPS, and 2.5 % DCM) was added to the resin with agitation for 2 hours protected from light. The resin was filtered and resulting solution was concentrated *in vacuo*. The peptide was triturated with cold diethyl ether and purified using reverse phase HPLC using $H_2O$/MeOH to yield **FI-LPMTG**. The sample was analyzed for purity using a Waters 1525 Binary HPLC Pump using a Phenomenex Luna 5u C8(2) 100A (250 x 4.60 mm) column; gradient elution with $H_2O/CH_3CN$.

**ESI-MS calculated $[M + H]^+$: , found:**
